# NucleicBERT: Deciphering the language of nucleic acids by a large-language model

**DOI:** 10.1101/2025.09.02.673754

**Authors:** Utkarsh Upadhyay, Julian Herold, Markus Götz, Alexander Schug

## Abstract

The vast majority of the human genome comprises non-protein-coding regions whose structural and functional roles remain poorly understood. Many of these regions function through RNA, yet progress in deep learning for RNA has lagged behind proteins because most methods rely on abundant structural labels or evolutionary alignments, both sparse for RNA. To address these challenges, we developed NucleicBERT, a self-supervised masked-language model that learns contextual representations capturing local and distal dependencies without requiring alignments or evolutionary information. Explainable AI analysis reveals that the model clusters RNA types in latent space and attends to structural properties like secondary structure and tertiary contacts, effectively “rediscovering” RNA biology from sequence correlations alone. When fine-tuned for downstream structural and functional tasks, NucleicBERT requires only single sequences, yet surpasses current state-of-the-art RNA models. This alignment-free framework addresses the scarcity of annotated 3D RNA data while providing a rapid, computational complement to experimental techniques. By bridging abundant unlabeled primary sequence corpora with more scarce structural annotations, NucleicBERT advances RNA structure prediction and provides insights into the working of LLMs. NucleicBERT is available at https://github.com/KIT-MBS/NucleicBERT.

## 1 Introduction

RNA plays essential roles in cellular processes ranging from transferring genetic information to forming the structural and catalytic core of the ribosome (1). Noncoding RNAs (ncRNAs) regulate gene expression (2; 3), yet the functionality of many remains unknown. This knowledge gap represents both a fundamental scientific challenge and an opportunity for therapeutic development. RNA-targeted drug design (4; 5) would particularly benefit from improved structure-function predictions, especially given that bacterial drug resistance (6) frequently involves ribosomal targets.

While RNA structure is critical for understanding function, experimental structure determination faces significant limitations. Although NMR spectroscopy and X-ray crystallography (7; 8) can provide detailed structural information, these approaches require substantial time and resources. Despite recent advances in cryo-EM (9), experimental bottlenecks persist, motivating the development of computational alternatives.

The success of deep learning in protein structure prediction (10; 11) has inspired similar efforts for RNA. However, RNA structure prediction faces a fundamental data scarcity problem: while sequencing technologies have generated millions of RNA sequences, corresponding structural data remains limited. This sequence-structure gap represents a major obstacle in computational structural biology.

Current computational approaches largely rely on evolutionary information extracted from multiple sequence alignments (MSAs). Direct Coupling Analysis (DCA) (12; 13; 14) pioneered the use of coevolutionary signals for protein structure prediction, with subsequent adaptation to RNA (15). Supervised methods such as RNAContact (16), CoCoNet (17) or the transformer-based model BARNACLE (18) build on these foundations. However, these approaches face practical limitations: evolutionary analysis is computationally expensive, and high-quality MSAs are unavailable for many RNA families.

Here, we present NucleicBERT, a large language model designed to overcome these limitations by learning directly from single RNA sequences. Trained on 30 million RNA sequences using masked language modeling, NucleicBERT achieves competitive or superior performance across diverse structural and functional prediction tasks without requiring evolutionary information. Based on the BERT architecture (19), our model treats nucleotides as tokens and RNA sequences as sentences, enabling it to capture rich contextual relationships through self-attention mechanisms (fig. 1, section 4.1).

**Figure 1:**
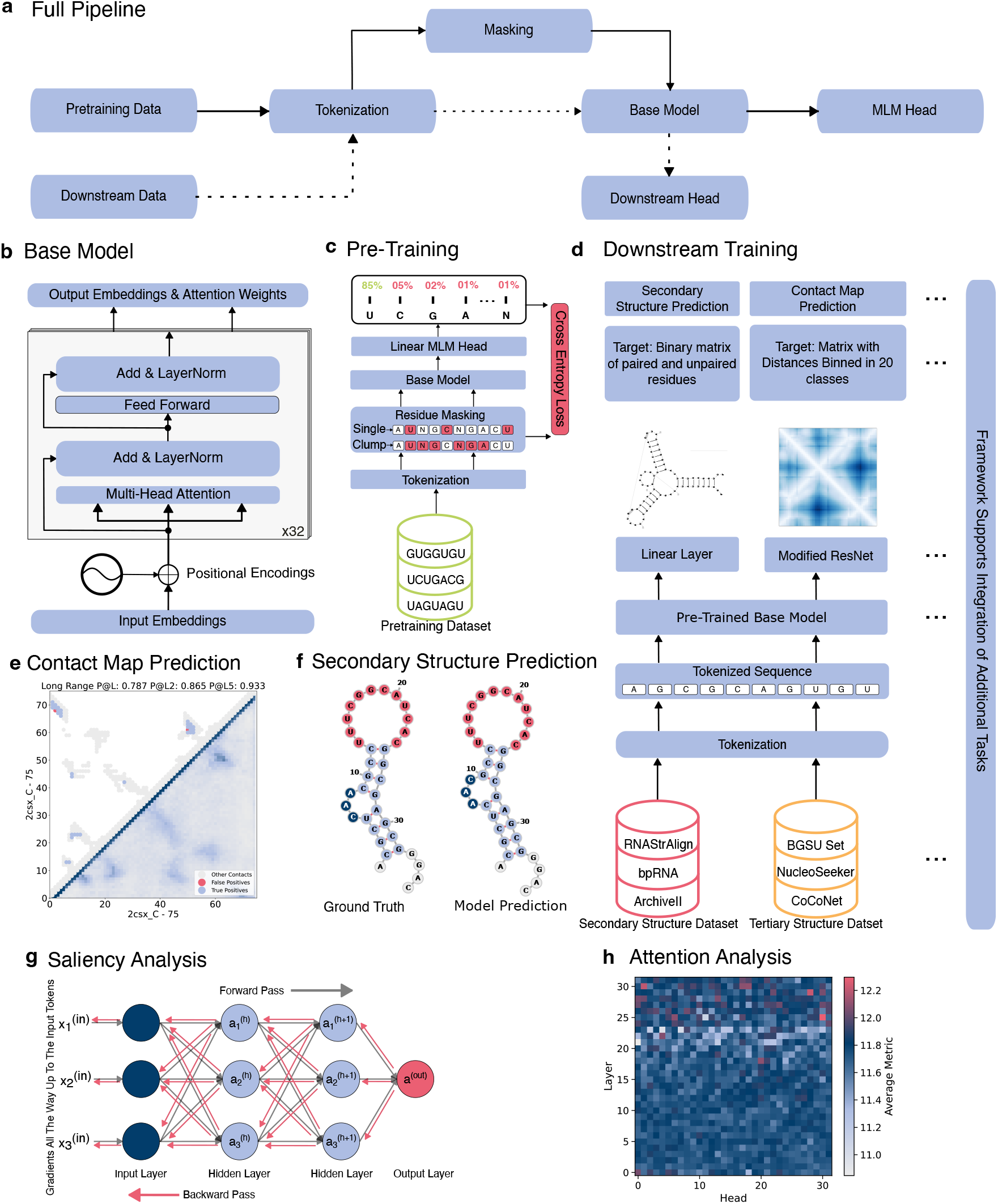
NucleicBERT architecture and training pipeline. **(a) Full pipeline** showing the complete workflow from data preprocessing through final predictions. **(b) Base model** architecture features transformer layers with multi-head attention and positional encodings to process sequence information. **(c) Pre-training** uses masked language modeling on large datasets to learn basic nucleic acid patterns and relationships.**(d) Downstream training** fine-tunes the pretrained model for specific tasks like secondary structure prediction and contact map prediction. We show only two tasks for brevity, and adding new tasks is straightforward. **(e-f) Task examples** shows sample results from our model. **(g-h) Model explainability** through saliency analysis reveals which input positions are most influential for predictions, while attention maps show how the model focuses on different sequence regions.

A critical challenge in applying deep learning to biological problems is model explainability. To address this, we conduct systematic explainable AI analysis to understand the biological knowledge NucleicBERT captures, an investigation never comprehensively applied to RNA language models. Through saliency analysis, attention weight examination, and perturbation study, we demonstrate how the model identifies structurally and functionally important sequence regions, providing insights into its decision-making process.

We make three primary contributions: (1) NucleicBERT, in its current form, achieves competitive or better performance compared to specialized tools across a range of structural and functional prediction tasks, (2) its modular architecture enables straightforward adaptation to new downstream applications, and (3) by incorporating explainable AI techniques, NucleicBERT goes beyond black-box modeling, enabling transparent interpretation of its learned representations. Together, these advances provide a versatile platform for RNA analysis that could accelerate both fundamental research and therapeutic development.

## 2 Results

Here, we present the results for NucleicBERT, demonstrate how pretraining leads to a deeper understanding of RNA sequences, describe the performance of our model on multiple downstream tasks, and analyze the explainability of this AI model.

### 2.1 Pretraining Performance Validation

To validate the effectiveness of our pretraining strategy, we conducted a comprehensive evaluation of the masked language modeling task on a subset of our validation data. NucleicBERT is trained to predict the identity of nucleotides that have been randomly masked out of RNA sequences:

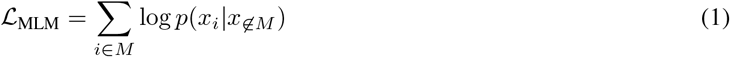

where for a randomly generated mask *M* that includes certain positions *i* in the sequence *x*, the model predicts the identity of the nucleotides *x*_*i*_ from the unmasked sequence *x*_∉*M*_ (section 4.1).

Our pretrained model achieved a mean accuracy of 0.831 on the masked token prediction task, representing a significant improvement over the non-pretrained model baseline (0.0708). This substantial performance gain demonstrates that our model captures complex sequence dependencies through the pretraining process (fig. 2a).

**Figure 2:**
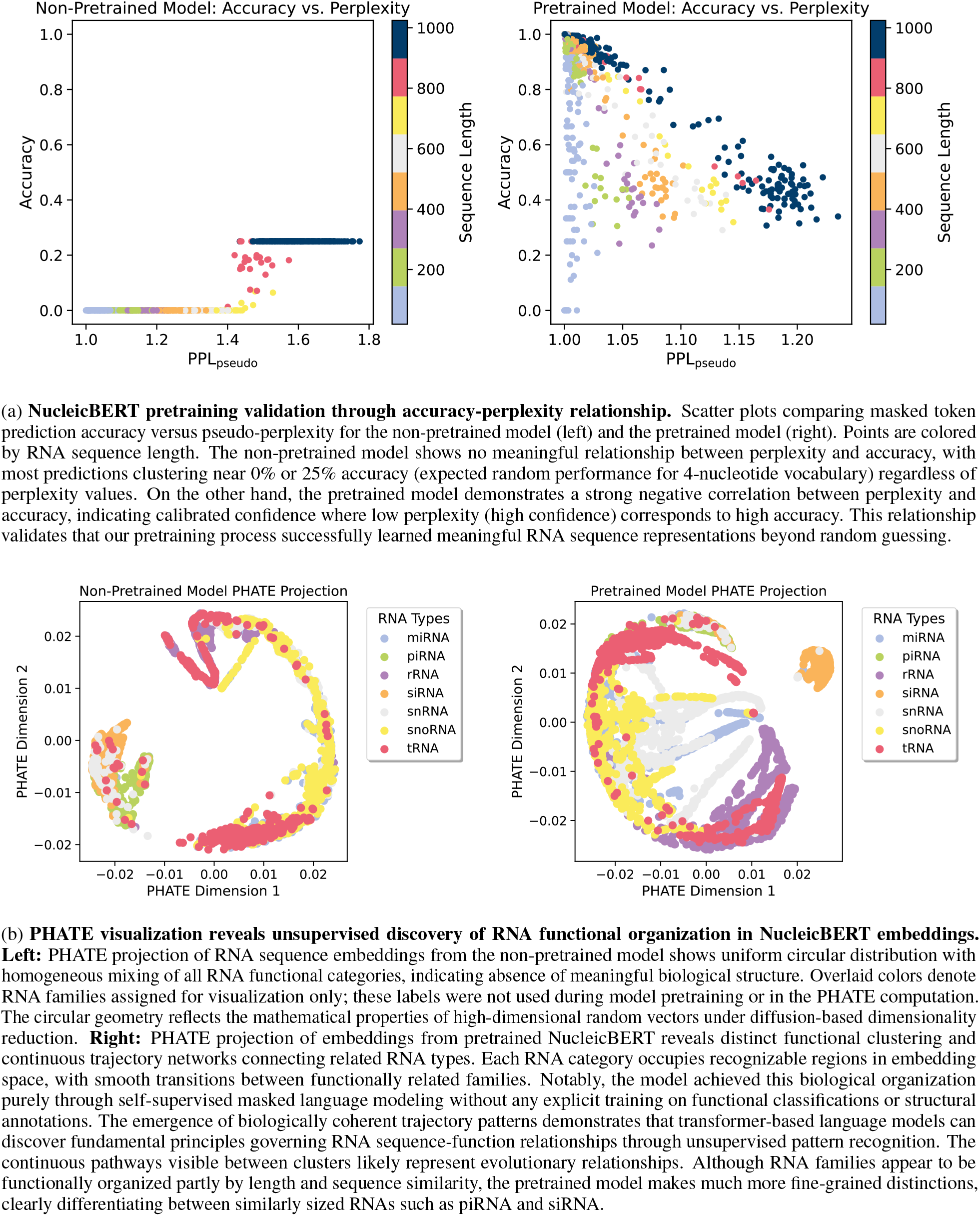
Pretraining validation and unsupervised biological discovery.

We evaluated model performance using perplexity, a standard metric for language models that measures the model’s uncertainty when predicting tokens. Perplexity is defined as the exponential of the negative log-likelihood of the sequence. For masked language models like NucleicBERT, we compute pseudo-perplexity (20) as:

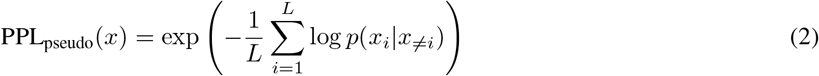

where *L* is the length of the input sequence and *p*(*x*_*i*_ | *x*_≠*i*_) represents the probability of nucleotide *x*_*i*_ given all other nucleotides in the sequence. Lower perplexity values indicate that the model confidently assigns high probabilities to correct predictions, while higher perplexity suggests greater uncertainty. Our model demonstrates a strong calibrated confidence relationship between perplexity and accuracy.

The high accuracy combined with the strong perplexity-accuracy correlation establishes that our model successfully learned to distinguish between probable and improbable RNA sequence patterns. These pretraining results provide empirical support for our approach and establish confidence in the quality of learned representations that we apply to subsequent structural prediction tasks and explainable AI analyses.

### 2.2 PHATE Analysis of RNA Sequence Embeddings

To further investigate the information captured by the embeddings of our model, we employed dimensionality reduction techniques. Our preferred method is PHATE (21) (Potential of Heat-diffusion for Affinity-based Trajectory Embedding), which addresses fundamental limitations of traditional dimensionality reduction methods when applied to biological data.

The comparative analysis (fig. 2b) between the non-pretrained and pretrained models provides definitive evidence of successful biological learning. Critically, NucleicBERT was never explicitly trained on RNA functional classifications; the model learned these patterns purely through self-supervised masked language modeling on sequence data.

The non-pretrained model produced a uniform circular distribution with mixing of RNA functional categories. A similar phenomenon was reported previously (22), where well-trained models show diverse and specialized representations while poorly trained models tend to have more homogeneous representations. Some observed clusters mostly arise from similarities in sequences and their lengths. In contrast, the pretrained model revealed complex trajectory networks with distinct functional clustering. Each RNA category formed recognizable regions connected by continuous pathways, demonstrating that the model discovered genuine sequence-function relationships through purely unsupervised learning.

The trajectory patterns reflect biologically plausible relationships between RNA types. Functionally related RNA families cluster together and connect through smooth transitions, suggesting that the model internalized evolutionary relationships and functional constraints purely from sequence context. Although a comprehensive analysis of these clustering patterns lies beyond the scope of the present work, preliminary observations reveal noteworthy trends. For instance, the similarly sized yet functionally distinct Piwi-interacting RNAs (piRNAs) and small interfering RNAs (siRNAs) are positioned adjacently in the non-pretrained model, whereas in the pretrained model, they are clearly separated.

### 2.3 Performance on Downstream Tasks

#### 2.3.1 Secondary Structure Prediction

To evaluate NucleicBERT’s capacity for secondary structure prediction, we conducted comprehensive benchmarking against established baseline methods using two widely adopted datasets in the RNA structure prediction community. Our evaluation employed the ArchiveII600 (33) dataset, a subset of ArchiveII containing 3,975 RNA structures from ten RNA families with sequence lengths under 600 base pairs, and the TS0 dataset from bpRNA-1m (34), comprising 1,305 test structures with lengths ranging from approximately 100 to 500 base pairs and 80% sequence identity clustering to ensure diversity (section 4.2).

Table 1a presents the comparison of NucleicBERT against several state-of-the-art methods, including traditional thermodynamic approaches (RNAfold (28)), recent deep learning models (MXfold2 (24), E2Efold (23)), and contemporary RNA language models (RNABERT (26), RNA-FM (25), RNAErnie+ (27)). Performance was assessed using precision, recall, and F1-score metrics calculated at the nucleotide level.

**Table 1:** NucleicBERT performance comparison across multiple downstream tasks.

(a) Secondary structure prediction performance
| Methods | ArchiveII600 |  |  | TS0 |  |  |
| --- | --- | --- | --- | --- | --- | --- |
| | Precision ↑ | Recall ↑ | F1 ↑ | Precision ↑ | Recall ↑ | F1 ↑ |
| E2Efold (23) | 0.738 | 0.665 | 0.690 | 0.140 | 0.129 | 0.130 |
| MXfold2 (24) | 0.788 | 0.760 | 0.768 | 0.519 | 0.646 | 0.558 |
| **NucleicBERT** | **0.915** | 0.848 | 0.872 | **0.719** | 0.610 | **0.649** |
| RNA-FM (25) | 0.752 | 0.737 | 0.744 | 0.518 | 0.620 | 0.564 |
| RNABERT (26) | 0.634 | 0.649 | 0.641 | 0.435 | 0.527 | 0.477 |
| RNAErnie+ (27) | 0.886 | **0.870** | **0.875** | 0.575 | **0.678** | 0.622 |
| RNAfold (28) | 0.565 | 0.627 | 0.592 | 0.494 | 0.631 | 0.536 |

(b) Contact map prediction performance
| Software | Top-L (L/5) ↑ | Top-L (L/2) ↑ | Top-L (L) ↑ |
| --- | --- | --- | --- |
| Non-Pretrained Model | 0.283 | 0.240 | 0.221 |
| **NucleicBERT** | **0.568** | **0.493** | **0.423** |
| RiNALMo (29) | 0.508 | 0.445 | 0.389 |
| RNA-FM (25) | 0.163 | 0.148 | 0.142 |

(c) Splice site prediction accuracy across species
| Model | Zebrafish ↑ | Fly ↑ | Worm ↑ | Plant ↑ | AVG ↑ |
| --- | --- | --- | --- | --- | --- |
| DNABERT (30) | 0.951 | 0.931 | 0.909 | 0.909 | 0.925 |
| **NucleicBERT** | 0.964 | **0.949** | **0.935** | **0.944** | **0.948** |
| RiNALMo (29) | **0.972** | 0.945 | 0.926 | 0.939 | 0.945 |
| SpliceBERT (31) | 0.957 | 0.946 | 0.934 | 0.936 | 0.943 |
| Spliceator (32) | 0.935 | 0.929 | 0.916 | 0.929 | 0.927 |

NucleicBERT demonstrates excellent performance across both evaluation datasets. On the ArchiveII600 benchmark, our model achieved comparable performance to state-of-the-art while using a simpler post-processing method for structure generation, see section 4.2.

On the more challenging TS0 dataset, NucleicBERT shows similar performance. The consistent performance advantage across diverse RNA families and sequence lengths demonstrates the robustness of the learned representations and their transferability to downstream structural prediction tasks.

#### 2.3.2 Distance and Contact Map Prediction

To assess NucleicBERT’s capability for distance and contact map prediction, with the latter resulting from thresholding the former, we conducted extensive evaluations on datasets derived from high-resolution RNA structures. Such a prediction represents a fundamental step toward understanding RNA tertiary structure, as it captures long-range nucleotide interactions and can act as a proxy for 3D structure prediction.

We evaluated NucleicBERT on two complementary datasets: a NucleoSeeker-generated (35) dataset containing 408 RNA structures and a BGSU (Bowling Green State University) (36) representative dataset comprising 467 structures (section 4.3).

In Table 1b, we compare NucleicBERT against current state-of-the-art RNA language models, including RiNALMo and RNA-FM. NucleicBERT outperformed all available methods. We calculate the top-*L* metrics for *≥* 24 residue separation, as this represents the most challenging aspect of this task, and these interactions are often critical for tertiary structure formation but cannot be inferred from local sequence patterns.

The importance of pre-training is also demonstrated in table 1b, where we also compare NucleicBERT’s pretrained model against a non-pretrained model. The pretrained model achieved remarkable improvements over the non-pretrained baseline. This substantial performance difference validates our hypothesis that masked language modeling on large-scale RNA sequence data enables the model to capture meaningful structural patterns that transfer effectively to other tasks.

#### 2.2.3 Splice Site Prediction

When genes are first copied from DNA to RNA, they contain both useful coding regions (exons) and non-coding segments (introns). RNA splicing removes non-coding segments (introns) and joins coding regions (exons) together to produce mature mRNA. Their precise prediction is critical for genome annotation and functional genomic studies.

To assess NucleicBERT’s capability in this functional prediction task, we evaluated its performance on splice site classification using established benchmark datasets (section 4.4).

Table 1c presents the comparison of NucleicBERT against established splice site prediction methods across four different species: zebrafish, fly, worm, and plant. The benchmark includes both traditional machine learning approaches (Spliceator (32)) and recent deep learning models (DNABERT (30), SpliceBERT (31), RiNALMo (29)).

The cross-species consistency in performance indicates that NucleicBERT has learned fundamental sequence-function relationships that are conserved across eukaryotic organisms. This capability positions the model as a valuable tool for genome annotation projects, particularly for newly sequenced or understudied species where species-specific training data may be limited.

#### 2.3.4 Fitness Prediction

To further assess NucleicBERT’s capacity for capturing RNA sequence-function relationships, we implemented a fitness prediction task using a comprehensive mutational dataset (37) of RNA sequences (section 4.6). Here, fitness refers to the self-cleavage activity of CPEB3 ribozyme variants. Specifically, fitness measures the proportion of RNA molecules that successfully cut their phosphodiester backbone. This measure provides a quantitative assessment of how sequence mutations affect catalytic efficiency.

To establish the value of pretrained representations, we compared NucleicBERT’s performance against a non-pretrained model. Since no results exist on this dataset in the literature for baseline comparison, we implemented a simple nearest neighbor-based estimator. This estimator averages the fitness values of all training sequences with a Hamming distance less than *k* = 3. While the non-pretrained model achieved a Pearson correlation coefficient of *r* = 0.67 between predicted and experimental fitness values, NucleicBERT reached *r* = 0.89, representing a 33% improvement in predictive accuracy. The enhanced performance was particularly pronounced for intermediate fitness ranges (0.2-0.8), where the null model exhibited poor calibration (fig. 3).

**Figure 3:**
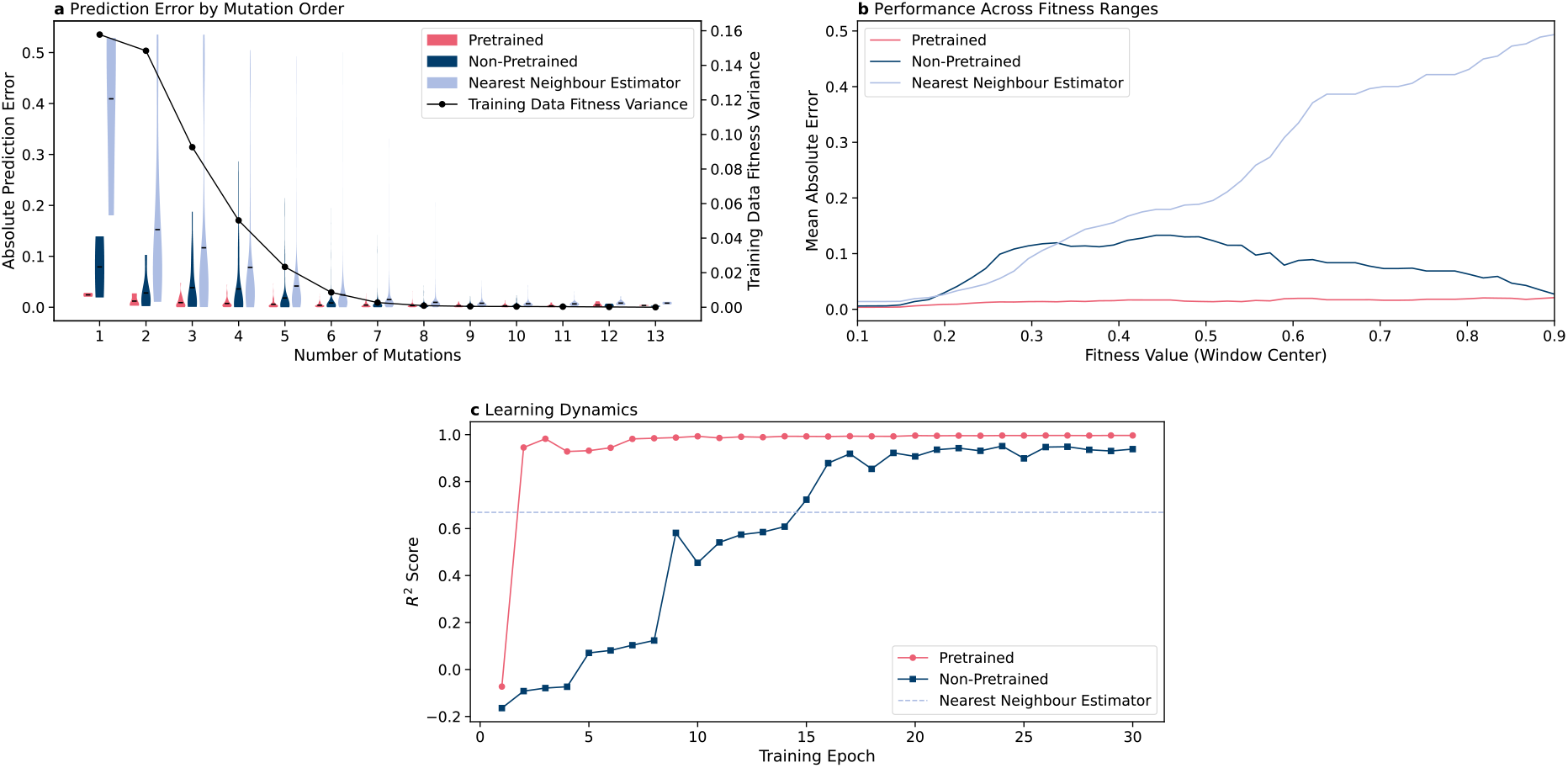
**a, Absolute prediction error** distributions across mutational complexity levels. The pretrained model (red) maintains consistently low prediction errors across all mutation counts, while the non-pretrained model (violet) exhibits the highest errors for sequences with few mutations and gradually improves with increasing mutational burden. The ball query baseline (blue) shows intermediate performance. All three models show good performance at higher order mutation values because the RNA sequences in the training data have extremely similar fitness values (black line, right y-axis) as mutational burden increases. **b, Mean absolute error** across fitness value ranges using sliding window analysis. The pretrained model maintains the lowest errors across the entire fitness spectrum, while the non-pretrained model shows elevated errors, particularly in intermediate fitness ranges. **c, Learning dynamics** showing *R*^2^ score progression during training. After some epochs, both models reach similar scores, but the low variance of fitness values at higher mutations allows easier prediction. The ball query baseline (horizontal line) represents non-learning similarity-based prediction.

Examining the prediction quality across mutational burden revealed additional insights into the model behavior. The pretrained model demonstrates (fig. 3) consistently excellent performance across all mutation values. In contrast, the non-pretrained model exhibits a different pattern; it performs worse on sequences with few mutations but improves as the number of mutations increases. The ball query baseline shows intermediate performance throughout. Notably, the training data fitness variance decreases sharply as mutation count increases, starting at 0.16 for single mutations and approaching zero for highly mutated sequences, suggesting that sequences with many mutations converge toward similar (likely low) fitness values. This pattern indicates that while individual mutations have highly variable fitness requiring sophisticated understanding to predict accurately, heavily mutated sequences become more predictable as they collectively drive fitness toward consistently poor outcomes.

### 2.4 Explainable AI Analysis

#### 2.4.1 Understanding Decision-Making through Saliency Analysis

We conducted a comprehensive saliency analysis by backpropagating loss gradients (38), (39) to identify input sequence positions that contribute most significantly to model predictions. This analysis was performed across three key tasks: masked token prediction, secondary structure prediction, and contact map prediction, revealing distinct patterns that provide insights into the model’s learned representations.

For the masked token prediction task (fig. 4a), saliency analysis confirmed that our model correctly focuses on the prediction objective (40), (41). Saliency values consistently showed pronounced spikes at positions containing `[MASK]`tokens, with the highest saliency scores occurring precisely at masked positions across sequences. This pattern validates that our masking strategy effectively directs the model’s focus to the prediction tokens while incorporating relevant local context, confirming that the pre-training objective is being optimized as intended and that the model has learned to utilize positional relationships for token prediction.

**Figure 4:**
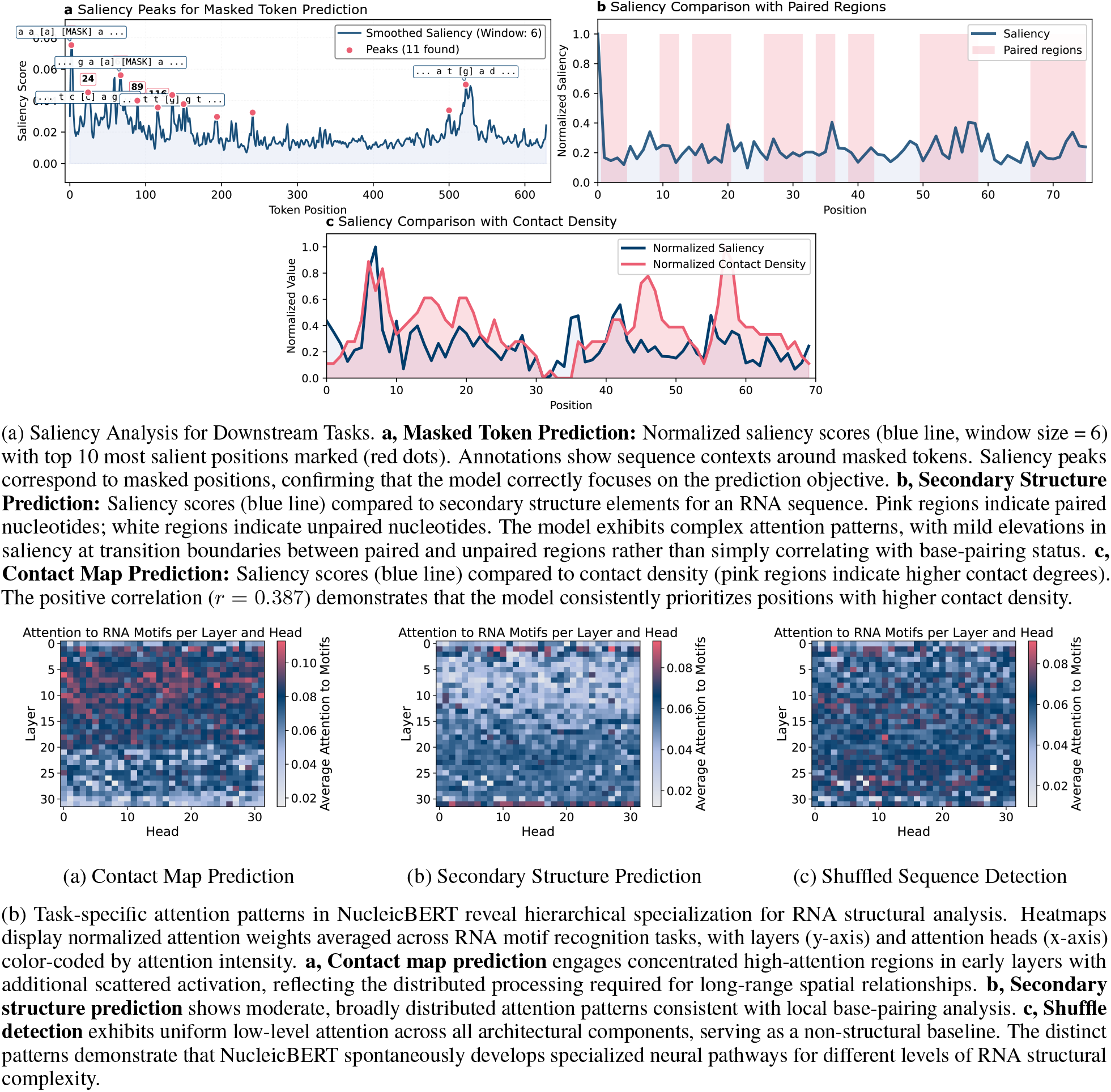
Explainable AI analysis of NucleicBERT.

For secondary structure prediction (fig. 4a), saliency analysis revealed a more complex relationship between the model’s focus and structural elements. While we observed only weak correlations between saliency and base-pairing status, the model showed heightened importance at structurally significant positions. Notably, the strongest saliency spikes occurred at transition points between paired and unpaired regions, suggesting that our model learned to identify structural boundaries and junction points that are critical for RNA folding (42).

The most compelling evidence for structural learning emerged from contact map prediction analysis (fig. 4a), where we observed significant positive correlations between contact density and saliency. Positions with higher contact degrees consistently received greater importance from the model. The correlation strength for contact prediction suggests that our model’s attention mechanisms are particularly well-suited for capturing long-range interactions and three-dimensional structural features. Collectively, these saliency analyses demonstrate that our RNA language model has developed a hierarchical understanding of RNA structure, from local sequence patterns during pre-training to complex structural parts in downstream tasks.

#### 2.4.2 Assessing Sequence Authenticity through Perturbation Sensitivity

Beyond understanding how the model makes decisions, we evaluated NucleicBERT’s capacity to distinguish authentic RNA sequences from artificially perturbed variants. This sequence authenticity assessment serves dual purposes: as a sensitivity analysis revealing the model’s learned constraints, and as a biologically meaningful downstream task for validating sequence integrity in real-world applications (section 4.5).

The model demonstrated remarkable sensitivity to sequence integrity, exhibiting a strong negative correlation between shuffle percentage and sequence authenticity classification (*r* = −0.951, *p* < 0.0001). Performance remained robust for minimally perturbed sequences, maintaining *>* 85% classification accuracy for shuffle levels below 10%. However, a pronounced performance decline occurred around the 20 − 25% shuffle threshold (fig. 5a).

**Figure 5:**
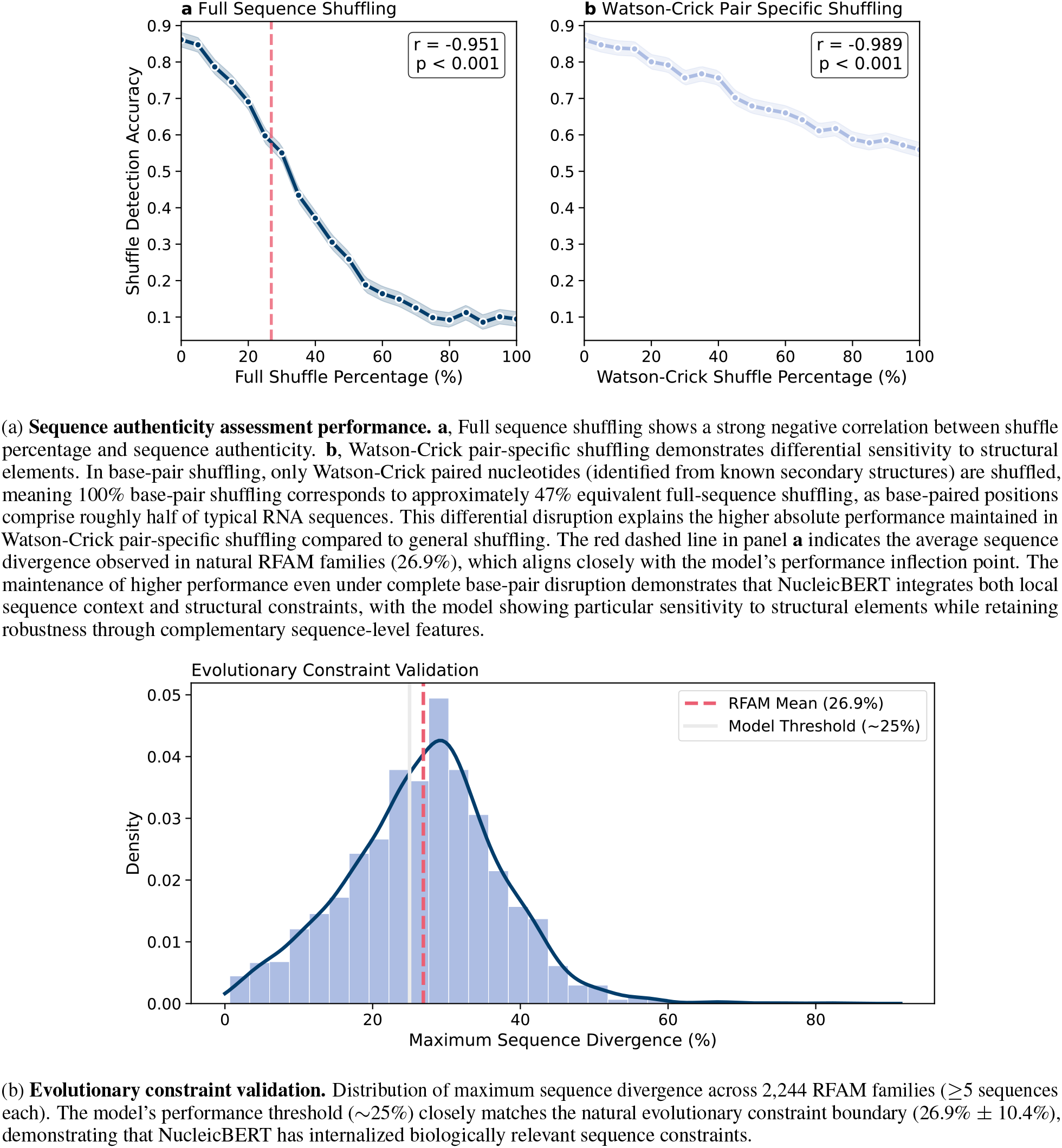
Biological validation of RNA sequence understanding through shuffle detection.

When perturbations were restricted to base-paired residues, the model exhibited an even stronger performance correlation (*r* = − 0.989, *p* < 0.0001), suggesting heightened sensitivity to disruptions in structurally important regions. Notably, even under complete base-pair shuffling (100%), the model maintained classification accuracy above 55%, indicating that it integrates both local sequence context and long-range structural constraints in its authenticity assessment (fig. 5a).

Natural RNA families tolerate an average sequence variation of 26.9% *±* 10.4% (section 4.5), closely matching our shuffle detection performance. The convergence between the model’s performance threshold (*∼* 25% shuffle) and the natural sequence variation boundary (26.9%) provides compelling evidence that NucleicBERT has internalized biologically relevant constraints on functional RNA sequence space (fig. 5b), while complementary work by Lambert et al. uses generative models to chart ribozyme diversity and functionality (43).

#### 2.4.3 Attention Patterns Reveal Structural Learning

To investigate how NucleicBERT processes different types of structural information, we analyzed normalized self-attention patterns across layers and heads for three distinct downstream tasks: contact map prediction, secondary structure prediction, and shuffle detection as a non-structural control task. Following established protocols for biological sequence transformers (44), we computed normalized attention weights to ensure meaningful cross-task comparison. The attention heatmaps reveal task-specific patterns that provide insights into the model’s internal representations of RNA structural complexity.

The three tasks demonstrate hierarchical attention specialization. Contact map prediction (fig. 4b) exhibits pronounced attention patterns, concentrated in early to middle layers (0–15) with scattered patterns in the deeper layers. This pattern suggests distributed processing of spatial relationships essential for tertiary structure formation (45). Secondary structure prediction (fig. 4b) shows heightened attention in the initial layers (1–2) with lower attention in the middle layers and the distributed pattern in the deeper layers. The deepest layer shows high attention again. (46). Shuffle detection (fig. 4b) exhibits a uniform attention pattern, providing a crucial baseline indicating minimal specialization for sequence-level discrimination.

The attention analysis reveals functional hierarchy within NucleicBERT’s architecture. Early layers (0–5) consistently show elevated attention across structural tasks, suggesting these components serve as general RNA feature extractors for local motifs and base-pairing patterns (47). Middle layers exhibit task-specific modulation: contact map prediction engages specialized high-attention regions while secondary structure prediction shows correspondingly lower attention in these same regions. The shuffle detection task lacks such specialized patterns, confirming that structural tasks engage distinct neural pathways. The deeper layers (21–29) show uniform attention patterns across all three tasks, suggesting these layers process sequence-level information rather than structural features.

The task-specific specialization mirrors RNA structural biology complexity: tertiary structure requires integration of multiple long-range interactions, secondary structure involves local base-pairing with contextual constraints, and sequence discrimination needs only general pattern recognition (48). The clear differentiation between tasks demonstrates that attention analysis, when properly normalized and validated against appropriate controls, provides valuable insights into RNA language model organization and their capacity for learning structural principles.

## 3 Discussion

NucleicBERT represents a significant advancement in RNA computational biology by achieving competitive performance across both structural and functional prediction tasks using only single-sequence inputs. Unlike previous specialized RNA tools that excel in specific domains, NucleicBERT demonstrates consistent performance across secondary structure prediction, contact map prediction, and splice site prediction, suggesting that our pretraining strategy captures fundamental principles governing RNA sequence-structure-function relationships. Additionally, new tasks can be added with modest computational requirements.

The elimination of multiple sequence alignment requirements addresses a critical limitation in the field. MSA-dependent methods face computational bottlenecks and limited applicability to novel RNAs or RNA families with shallow MSAs. Our single-sequence approach maintains competitive accuracy while offering practical advantages.

Our comprehensive explainable AI analysis reveals that NucleicBERT spontaneously develops task-specific specialization that mirrors RNA structural complexity. The attention patterns demonstrate architectural components with distinct roles. The saliency analysis confirms biological learning through heightened sensitivity to structurally critical regions. The biological validation through shuffle detection provides compelling evidence that these capabilities reflect genuine understanding rather than statistical artifacts. The remarkable convergence between our model’s performance threshold (*∼* 25% shuffle) and the natural sequence variation boundary observed in Rfam families (26.9%) indicates that NucleicBERT has internalized biologically relevant constraints. This biological grounding distinguishes our approach from other methods.

A significant finding is the unsupervised discovery of biological organization revealed through PHATE analysis. The model’s clustering of RNA sequences by functional categories, without explicit functional annotations during pretraining, suggests that sequence-function relationships can be discovered through purely self-supervised learning. This capability positions NucleicBERT as a powerful tool for annotating the many non-coding genomic regions that remain functionally uncharacterized.

NucleicBERT delivers competitive or superior performance compared with other methods across diverse RNA tasks, establishing it as a unifying framework with implications for, e.g., RNA-targeted drug discovery. The model’s explainability features enable integration with experimental workflows, where predictions can guide hypothesis formation and experimental design. As high-throughput techniques for measuring RNA function continue to expand, NucleicBERT’s pretrained representations provide a computational foundation for accelerating sequence-function relationship discovery across the transcriptome, bridging abundant sequence data with scarce functional annotations to advance RNA biology and therapeutic applications.

## 4 Methods

### 4.1 Pre-training Architecture and Dataset

Our model consists of 32 transformer layers, each with 32 attention heads and an embedding dimension of 1,024. We employed learned positional encodings to capture position-specific information.

For tokenization, we trained a Byte Pair Encoding (BPE) tokenizer with a clump size of 1, effectively creating a character-level tokenization strategy in which each nucleotide or ambiguity character becomes its own token. This approach maintains maximal granularity without any merge operations, which is well-suited to the relatively small RNA alphabet.

The resulting vocabulary size is 25. It comprises the four standard RNA nucleotides (A, U, G, C), sixteen ambiguity characters, and five special tokens. The ambiguity codes extend beyond conventional IUPAC representations and were empirically determined by identifying all unique non-standard characters present in our training dataset. The special tokens serve distinct roles: `[PAD]`is used for padding sequences within a batch; `[UNK]`denotes unknown or out-of-vocabulary characters; `[MASK]`is used during masked language modeling; `[CLS]`is prepended to each sequence to aggregate global context, and `[SEP]`separates input segments when required for downstream tasks.

The model was pretrained using a masking strategy adapted from BERT-style language modeling (19), combining both token-level and subsequence-level masking to encourage learning of local and higher-order sequence patterns. For token-level masking, 15% of tokens in each input sequence were selected at random. Of these, 80% were replaced with the `[MASK]` token, 10% were substituted with a random token from the vocabulary, and 10% were left unchanged. In parallel, a subsequence masking strategy was employed, in which contiguous spans of 4 to 8 tokens were randomly selected and masked as a group. This span-based approach draws on principles from SpanBERT (49) and is intended to promote learning of biologically meaningful RNA motifs, which often appear as short subsequences. We also evaluated motif-based masking as suggested by RNAErnie (27), but did not observe any downstream performance improvement.

To train NucleicBERT, we obtained non-coding RNA sequences from the MARS database (50). It contains *∼*1.7 billion sequences in FASTA format, but since we wanted to train our model only on non-coding RNA sequences, we extracted all sequences with the `ncRNA` keyword in their description and obtained *∼*30 million sequences. 80% of this data was used in training, while the remaining was used in validation. We trained this 404-million-parameter model using 192 A100-40-GB GPUs with a linear learning rate schedule and a batch size of 16. To ensure stable gradients and prevent overfitting, we maintained a dropout rate of 0.1 and clipped gradient values to 0.001. Each epoch of training required approximately 1 hour and 20 minutes, and the model was trained for 300 epochs. Pre-training accuracy plateaued at 83.1% on the validation dataset.

For analysing embeddings obtained from our model, we used the PHATE dimensionality reduction technique. Unlike clustering-focused approaches such as t-SNE (51) or PCA (52), PHATE preserves continuous trajectories and branching structures through a diffusion-based framework that simulates random walks across data manifolds to capture global relationships.

PHATE constructs embeddings through a multi-step process: computing local similarities with *α*-decaying kernels, creating Markovian transition matrices, and raising these matrices to successive powers to simulate longer diffusion processes. This diffusion operation denoises data while learning global relationships between distant points. Rather than using diffusion coordinates directly, PHATE computes an information-geometric “potential distance” between probability distributions, making it particularly effective at preserving biological transitions.

### 4.2 Secondary Structure Architecture and Dataset

For secondary structure prediction, we developed a binary classification framework that determines whether nucleotide pairs form base-pairing interactions. The approach involves attaching a bottleneck residual network architecture to the pretrained NucleicBERT model, which processes pairwise nucleotide representations generated through outer concatenation of the transformer embeddings. Specifically, the representation for nucleotide pair (*i, j*) is constructed by concatenating the embedding vectors of positions *i* and *j* from NucleicBERT’s final layer.

The prediction network generates a symmetric probability matrix indicating base-pairing likelihood for each nucleotide pair in the sequence. We optimized the model using a binary cross-entropy loss, computing gradients exclusively for the upper triangular portion of the matrix to account for base-pairing symmetry constraints. Training proceeded for 50 epochs using a learning rate of 1 × 10^−5^, and we trained all model parameters simultaneously without implementing gradual unfreezing strategies.

We converted the predicted probability matrix into discrete secondary structures using a greedy decoding algorithm. This iterative process selects the highest-probability nucleotide pair as base-paired, then removes conflicting pairs from consideration in subsequent iterations. The algorithm enforces biological constraints by excluding non-Watson-Crick pairings and preventing formation of hairpin loops shorter than four nucleotides (constraint: |*i* − *j*| *≥* 4). We determined the optimal probability threshold through validation set optimization to achieve balanced pairing ratios.

Performance evaluation incorporated RNA structural flexibility by accepting near-correct predictions as valid. Following established evaluation protocols (53), we considered predictions within one position of the true pairing as correct: for a reference pair (*i, j*), the predictions (*i ±* 1, *j*) and (*i, j ±* 1) were scored as accurate. Final F1 scores represent the average of individual structure-level F1 calculations across the entire test set. Several benchmark datasets are used in this study:

1. The RNAStrAlign (54) dataset comprises 37,149 RNA structures from eight distinct RNA families, with sequence lengths ranging from approximately 100 to 3,000 base pairs.
2. ArchiveII (33) dataset consists of 3,975 RNA structures from ten RNA families, with lengths ranging from approximately 100 to 2,000 bp. We apply a cutoff of 600 bp to get the standard dataset used in tools like UFold and RNAErnie.
3. bpRNA-1m (34), contains a large number of highly similar sequences, thus a sequence identity cutoff of 80% is applied. The dataset is randomly split into three subsets: TR0 (10,814 structures) for training, TV0 (1,300 structures) for validation, and TS0 (1,305 structures) for testing.

The model was trained on the RNAStrAlign data and TR0 data combined. We tested the model on ArchiveII600 and TS0 datasets.

### 4.3 Tertiary Structure Prediction Architecture and Dataset

To use the representations learned during the pre-training, we finetuned our model to predict distance maps and contact maps for a given RNA sequence.

Distance maps represent spatial pairwise inter-residue interactions, providing an RNA’s translationally and rotationally invariant topological representation. For a sequence of length *L*, a distance map is a matrix (*L* × *L*) representation that delineates pairwise interactions between atoms within a bio-molecular structure. The elements of the matrix correspond to specific residues, and the entry at the intersection of row *i* and column *j* indicates the Euclidean distance between them.

Contact maps are a binary representation derived from these distance maps, where a threshold distance is applied to determine whether two nucleotides are in contact. If the distance between nucleotides is below the defined threshold (we used 8 Å(55)), they are considered to be in contact (represented as 1 or a marked point), while distances above the threshold are considered non-contacts (represented as 0 or empty space). This simplified binary representation captures essential structural information while reducing noise and computational complexity.

The PDB database contains merely 8,446 (as of January 2025) annotated tertiary RNA structures, reflecting the scarcity of such data. Further, a large proportion of these structures are hybrids, have poor resolutions, have small lengths, and have high sequence similarity. An unfiltered dataset with redundancy, obtained from the PDB database, can impact the generalizability of the model. We used the BGSU RNA 3D representative dataset (release 3.368, January 2025 (36)) as our starting point. We processed this dataset by filtering for structures with lengths between 32-1,024 nucleotides, removing redundancy using CD-HIT-EST (56) with an 80% sequence identity threshold, and ensuring structural quality. For each selected structure, we extracted the sequence, resolution, and chain information from mmCIF files, creating a curated dataset suitable for model training. We also used NucleoSeeker (35) to create a non-redundant dataset for downstream training using the same steps as described above. Both datasets were used separately for training, and they showed similar performance.

To prepare distance maps, the distances between the heavy atoms of each residue are calculated with every other residue. These distances are assigned into 20 classes such that:

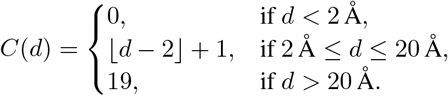

Many structures in the PDB database have missing residues, which can cause problems in the training because there will be a mismatch in the length of sequences and the shape of the distance map. To solve this, we use PDBFixer (57) to add missing residues and atoms. Further, we use the Chemical Component Dictionary from PDB to replace the remaining non-standard residues.

Lastly, we remove the trivial distances from the distance maps because they originate from the backbone and do not contain significant information. We remove four off-diagonals above and below the main diagonal, including the main diagonal. Data prepared in this manner allows the model to learn long-range interactions.

For the downstream supervised model, we use image segmentation techniques. The attention weights representation from the pretrained model is fed into a ResNet (58) model, which classifies each pixel (element) of the attention weights into the 20 categories.

Contact map prediction presents unique challenges that differentiate it from conventional computer vision tasks. While distance maps can be interpreted as 2D images—making contact prediction superficially analogous to pixel-level image labeling—this analogy is limited and should be applied with caution. Some techniques from image labeling may be transferable, but there are several important distinctions.

First, in typical image classification or segmentation tasks, input images are often resized to a fixed dimension. In contrast, contact maps must preserve the native dimensions of the RNA sequence, as predictions are required for every pair of residues. Second, the input features for contact prediction are substantially more complex, incorporating both sequential and pairwise features, unlike the raw pixel intensities used in standard image tasks. Third, the label distribution in contact maps is highly imbalanced: the proportion of positive contacts is typically less than 2%, creating a significant challenge for model training (59).

### 4.4 Splice Site Prediction Architecture and Dataset

We formulated splice site prediction as a binary classification problem, where the model must distinguish between authentic splice sites and non-functional sequences. The evaluation was conducted using carefully curated multi-species datasets that have become standard benchmarks in the community.

For splice site prediction evaluation, we employed the benchmark datasets established in the Spliceator (32) study, which have since been adopted by the RiNALMo framework (29). These datasets represent a gold standard for multi-species splice site prediction assessment due to their rigorous curation and phylogenetic diversity.

The benchmark datasets encompass four evolutionarily diverse species: zebrafish (*Danio rerio*), fruit fly (*Drosophila melanogaster*), nematode worm (*Caenorhabditis elegans*), and thale cress (*Arabidopsis thaliana*). Each benchmark contains 10,000 splice site sequences and 10,000 non-splice site sequences, maintaining a balanced 1:1 ratio to prevent class imbalance bias. The datasets include both canonical and non-canonical splice sites, with non-canonical splice sites representing: 307 in human, 85 in zebrafish, 120 in fly, 67 in worm, and 122 in plant samples.

The original Spliceator datasets were constructed from high-quality genomic sequences obtained from the Ensembl database release 87, with splice sites extracted and flanked by ±300 nucleotide environments to provide sufficient sequence context. A verification process ensured that no sequences containing undetermined nucleotides (noted as ‘N’) were included in the datasets. Each sequence spans 600 nucleotides total, with the GT (donor) or AG (acceptor) dinucleotide positioned at the central location (positions 301-302).

The negative subset construction employed a heterogeneous approach, incorporating three categories of non-splice site sequences: randomly selected exon regions, randomly selected intron regions, and false positive sequences containing GT or AG dinucleotides that do not correspond to authentic splice sites. This heterogeneous negative dataset design (termed GS_1 in the Spliceator methodology) was shown to achieve superior performance compared to homogeneous negative datasets containing only false positive sequences.

The benchmark datasets underwent rigorous quality control procedures based on the G3PO+ multi-species benchmark methodology. Sequences were classified into ‘Confirmed’ (error-free) and ‘Unconfirmed’ (containing potential gene prediction errors) categories through expert-guided comparative sequence analysis. Only ‘Confirmed’ sequences were retained in the final benchmarks to ensure biological authenticity of the splice sites.

To prevent sequence redundancy and potential overfitting, duplicate sequences were systematically removed from each dataset. Sequence similarity analysis revealed that the majority (>90%) of sequences in each benchmark share between 20-30% identity for full-length sequences, with higher conservation (20-60% identity) observed in the immediate splice site vicinity.

For this downstream task, we fine-tuned NucleicBERT using a 2-layer sequence-level classification head that processes the model’s contextual embeddings to make binary predictions. The fine-tuning process was performed separately for donor and acceptor splice sites to optimize performance for each specific recognition task, following established protocols from both the Spliceator and RiNALMo methodologies.

### 4.5 Shuffled Sequence Detection Architecture and Dataset

We assessed NucleicBERT’s capacity to identify authentic RNA sequences by systematically introducing controlled perturbations through nucleotide shuffling. Using the same dataset employed for secondary structure prediction training and testing, we generated shuffled variants with perturbation levels ranging from 0% to 100% in 5% increments. At each shuffle percentage, we measured the fraction of sequences that the model still classified as authentic RNA. In this study, we use the embeddings of the pretrained model, which are passed through a 3-layer perceptron to predict if a given sequence is shuffled or not.

To investigate whether the model preferentially attends to structurally critical regions, we performed targeted perturbation analysis focusing exclusively on Watson-Crick base-paired positions identified from known secondary structures. This approach allowed us to assess the model’s understanding of the hierarchical constraints that govern RNA folding.

To establish whether the observed performance thresholds reflect genuine biological constraints rather than training artifacts, we analyzed sequence variation patterns across natural RNA families. We extracted seed multiple sequence alignments for 4,178 RNA families from the Rfam database (60) (release 15.0), focusing on families containing at least five sequences to ensure statistical reliability. For each family, we identified a representative sequence by optimizing Levenshtein similarity across all family members and calculated the maximum sequence variation tolerance within each family using normalized edit distance.

### 4.6 Fitness Prediction Architecture and Dataset

We use the CPEB3 mutation dataset (37), which represents a comprehensive mutagenesis study of RNA sequences that serve as substrates for CPEB3 (Cytoplasmic Polyadenylation Element-Binding Protein 3), an RNA-binding protein that regulates gene expression through control of mRNA polyadenylation. This dataset contains fitness landscape data from a high-throughput cleavage assay where systematically mutagenized RNA sequences were evaluated for their susceptibility to cleavage by CPEB3. Each RNA variant was tested across three independent biological replicates, with fitness calculated as the ratio of cleaved to uncleaved RNA molecules, providing a quantitative measure of how effectively each sequence serves as a CPEB3 substrate. The rigorous experimental methodology, including multiple biological replicates and standardized cleavage assays, ensures high-quality fitness measurements suitable for downstream computational analysis and machine learning models designed to predict RNA substrate fitness from sequence information.

For fitness prediction, we employed a task-specific regression head that processes the pooled embeddings from NucleicBERT’s final transformer layer. The regression architecture consists of a 3-layer perceptron that maps the 1,024-dimensional RNA sequence representations to scalar fitness predictions. We implemented an 80/20 train-test split, ensuring that the evaluation dataset contained sequences spanning the full mutational spectrum while maintaining statistical power for performance assessment.

To establish a biologically informed baseline that captures the assumption that similar fitness values arise from sequences with similar mutation patterns, we develop a ball-query model, which is essentially a k-nearest neighbors model. It measures the distance between sequences using the symmetric distance between mutation sets, and for each target sequence, it finds all reference sequences within a specified cutoff radius. The fitness value is the average fitness of all sequences within the radius.

## Acknowledgements

The authors gratefully acknowledge the Gauss Centre for Supercomputing e.V. (www.gauss-centre.eu) for funding this project by providing compute time through the John von Neumann Institute for Computing (NIC) on the GCS Supercomputer JUWELS (61) at Jülich Supercomputing Centre (JSC). Additional computing time was provided on the supercomputer JURECA (62) at Forschungszentrum Jülich under grant EatsRNA. A.S. and J.H. recognize support by HIDSS4Health – the Helmholtz Information & Data Science School for Health. M.G., A.S., and J.H. recognize support by the Helmholtz Association’s Initiative and Networking Fund (INF) under the Helmholtz AI platform grant and the grant ZebraTwin. A.S. recognizes support by the Helmholtz Foundation Model Initiative (HFMI) of the Helmholtz Association under the grants PROFOUND and Virtual Cell. The funders had no role in study design, data collection, analysis, decision to publish, or preparation of the manuscript. The authors thank Stefan Kesselheim (JSC/FZJ), Jiangtao Wang (JSC/FZJ), Christian Faber (JSC/FZJ), and Anton Dorn (JSC/FZJ) for valuable discussions.

## Data Availability

### Pre-training Data

The RNA sequences used for pre-training were obtained from the publicly available MARS database (50), specifically extracting *∼*30 million non-coding RNA sequences with the ncRNA keyword.

### Benchmark Datasets

All evaluation datasets used in this study are publicly available:

- Secondary structure prediction: ArchiveII (33), bpRNA-1m (34), and RNAStrAlign (54).
- Tertiary structure prediction: PDB structures processed through NucleoSeeker (35) and BGSU RNA 3D representative dataset (36) (release 3.368, January 2025).
- Splice site prediction: Spliceator (32) benchmark datasets for zebrafish, fly, worm, and plant species.
- Fitness prediction: CPEB3 mutation dataset as (37)
- Shuffle detection: ArchiveII (33), bpRNA-1m (34), RNAStrAlign (54), and Rfam database release 15.0 (60).

Model weights are available at https://doi.org/10.5281/zenodo.16989562. Further processed datasets generated for this study are available upon reasonable request from the corresponding author.

## Contributions

**Conceptualization**: U.U. and A.S. conceived and designed the study. **Software:** U.U. developed the main software package. J.H. and M.G. contributed code for the explainable AI components. **Methodology:** U.U., J.H. and M.G. provided the analytical framework and method validation. **Formal analysis:** All authors contributed to the formal analysis. **Writing – original draft:** All authors contributed to writing the original manuscript. **Writing – review & editing:** All authors reviewed and approved the final manuscript.

