## Supplementary Information for "NucleicBERT: Deciphering the language of nucleic acids by a large-language model"

**Utkarsh Upadhyay**
Jülich Supercomputing Centre
Forschungszentrum Jülich
52428 Jülich


**Julian Herold**
Scientific Computing Centre
Karlsruhe Institute of Technology
Karlsruhe


**Markus Götz**
Helmholtz AI
Scientific Computing Centre
Karlsruhe Institute of Technology
Karlsruhe


**Alexander Schug** *
Jülich Supercomputing Centre
Forschungszentrum Jülich
Jülich


#### Overview

This document presents the complete hyperparameter configurations for NucleicBERT. We provide the parameters used in NucleoSeeker for curating the tertiary structure dataset. In addition, we cover the hyperparameters used for pretraining and various downstream tasks, including secondary structure prediction, splice site prediction, shuffle detection, fitness prediction, and contact map/distance map prediction. These hyperparameters enable reproducible implementation of the NucleicBERT framework for RNA structural and functional prediction tasks.

#### 1 NucleoSeeker Dataset Generation Parameters

Table 1: NucleoSeeker dataset generation parameters used for RNA structure data collection from the RCSB PDB database.

| **Parameter** | **Value** |
|---|---|
| Structure Determination Methodology | experimental |
| RCSB Entity Polymer Type | RNA |
| Polymer Type | polyribonucleotide |
| Resolution (Å) | 4.0 |
| Year Range | 2025 |
| Sequence Length Range | (32, 1024) |
| Sequence Identity (%) | 80.0 |
| E-value (CMScan) | 10 |
| Alignment Tool | clustal-omega |

*Corresponding author

#### 2 NucleicBERT Pre-Training on RNA Sequences

Table 2: NucleicBERT Pre-Training Hyperparameters

| Parameter | Value |
|---|---|
| **Model Architecture** | |
| Dropout rate | 0.1 |
| Embedding dimension | 1024 |
| Maximum sequence length | 1024 |
| Number of attention heads | 32 |
| Number of hidden layers | 32 |
| Positional encodings | Learned |
| Vocabulary size | 25 |
| **Training Configuration** | |
| Batch size per device | 16 |
| Gradient accumulation steps | 4 |
| Gradient clipping algorithm | Value-based |
| Gradient clipping value | 0.001 |
| Ignore index | -100 |
| Learning rate | $1.0 \times 10^{-5}$ |
| Learning rate scheduler | Linear with warmup |
| Loss function | Cross-entropy |
| Mask probability | 0.15 |
| Optimizer | AdamW |
| **Data Configuration** | |
| Data split ratio | 0.8/0.2 |
| **Distributed Training** | |
| Accelerator | GPU |
| Devices per node | 4 |
| Number of nodes | 48 |
| Number of workers | 32 |
| Precision | 16-bit mixed |
| Strategy | DDP |

#### 3 Secondary Structure Prediction Fine-tuning

Table 3: Secondary Structure Prediction Model Hyperparameters

| Parameter | Value |
|---|---|
| **Model Architecture** | |
| Base model | Pre-Trained NucleicBERT |
| Downstream Architecture | ResNet2D + Convolution |
| Downstream Input | Embeddings Outer Product |
| Kernel size | 3 |
| Normalization | InstanceNorm2D |
| Number of ResNet blocks | 2 |
| Output channels | 1 |
| **Training Configuration** | |
| Learning rate | $1.0 \times 10^{-5}$ |
| Learning rate scheduler | Cosine with warmup |
| Loss function | BCEWithLogitsLoss |
| Optimizer | AdamW |
| Warmup steps | 10% of max epochs |
| **Data Configuration** | |
| Maximum sequence length | 1000 |
| Minimum sequence length | 0 |
| **Distributed Training** | |
| Accelerator | GPU |
| Number of devices | 4 |
| Number of workers | 32 |
| Precision | 16-bit mixed |
| Strategy | DDP |

### 4 Splice Site Prediction Fine-tuning

Table 4: Splice Site Prediction Model Hyperparameters

| **Parameter** | **Value** |
|---|---|
| **Model Architecture** | |
| Base model | Pre-Trained NucleicBERT |
| Downstream Architecture | Linear(1024→128) + GELU + Linear(128→1) |
| Downstream Input | [CLS] Token Representation |
| **Training Configuration** | |
| Batch size per device | 4 |
| Learning rate | $1.0 \times 10^{-6}$ |
| Learning rate scheduler | Cosine with warmup |
| Loss function | BCEWithLogitsLoss |
| Maximum epochs | 30 |
| Optimizer | AdamW |
| Warmup steps | 10% of max epochs |
| **Data Configuration** | |
| Data split ratio | 0.8/0.2 |
| Maximum sequence length | 400 |
| Minimum sequence length | 0 |
| **Distributed Training** | |
| Accelerator | GPU |
| Number of devices | 4 |
| Number of nodes | 1 |
| Number of workers | 24 |
| Strategy | DDP |

#### 5 Shuffle Detection Fine-tuning

Table 5: Shuffle Detection Model Hyperparameters

| Parameter | Value |
|---|---|
| **Model Architecture** | |
| Base model | Pre-Trained NucleicBERT |
| Downstream Architecture | Linear(1024→256) + ReLU + Dropout(0.2) + Linear(256→64) + ReLU + Dropout(0.2) + Linear(64→1) |
| Downstream Input | Final Layer Embeddings |
| Attention pooling | Linear(1024→256) + Tanh + Linear(256→1) |
| **Training Configuration** | |
| Batch size per device | 4 |
| Learning rate | $5.0 \times 10^{-5}$ |
| Learning rate scheduler | Cosine with warmup |
| Loss function | BCEWithLogitsLoss |
| Optimizer | AdamW |
| Warmup steps | 10% of max epochs |
| **Data Configuration** | |
| Data split ratio | 0.8/0.2 |
| Maximum sequence length | 1024 |
| Minimum sequence length | 20 |
| **Distributed Training** | |
| Accelerator | GPU |
| Number of devices | 4 |
| Number of nodes | 1 |
| Number of workers | 24 |
| Strategy | DDP |

#### 6 Fitness Prediction Fine-tuning

Table 6: Fitness Prediction Model Hyperparameters

| Parameter | Value |
|---|---|
| **Model Architecture** | |
| Base model | Pre-Trained NucleicBERT |
| Downstream Architecture | Linear(1024→128) + ReLU + Dropout(0.1) + Linear(128→64) + ReLU + Dropout(0.1) + Linear(64→1) |
| Downstream Input | [CLS] Token Representation |
| **Training Configuration** | |
| Batch size per device | 4 |
| Learning rate | $1.0 \times 10^{-6}$ |
| Learning rate scheduler | Cosine with warmup |
| Loss function | L1Loss |
| Optimizer | AdamW |
| Warmup steps | 10% of max epochs |
| **Data Configuration** | |
| Data split ratio | 0.8/0.2 |
| Maximum sequence length | 1024 |
| Minimum sequence length | 20 |
| **Distributed Training** | |
| Accelerator | GPU |
| Number of devices | 4 |
| Number of nodes | 1 |
| Number of workers | 24 |
| Strategy | DDP |

### 7 Contact Map and Distance Map Prediction

Table 7: Contact Map and Distance Map Prediction Model Hyperparameters

| **Parameter** | **Value** |
|---|---|
| **Model Architecture** | |
| Base model | Pre-Trained NucleicBERT |
| Downstream Architecture | ResidualNetwork2D |
| Downstream Input | Attention weights |
| Frozen backbone | False |
| Input channels | 1024 (32 heads × 32 layers) |
| Number of residual blocks | 1 |
| Output channels (contact map) | 2 |
| Output channels (distance map) | 20/21 |
| **Training Configuration** | |
| Batch size per device | 1 |
| Gradient clipping algorithm | Norm-based |
| Gradient clipping value | 1.0 |
| Learning rate | $1.0 \times 10^{-6}$ |
| Learning rate scheduler | Cosine with warmup |
| Optimizer | AdamW |
| Warmup steps | 10% of max epochs |
| **Loss Configuration** | |
| Distance threshold | 8.0 Å |
| Focal loss alpha | 0.8 |
| Focal loss gamma | 2.0 |
| Ignore index (contact map) | -1 |
| Ignore index (distance map) | 21 |
| Loss function | Focal loss |
| **Data Configuration** | |
| Data split ratio | 0.8/0.2 |
| Maximum sequence length | 512 |
| Minimum sequence length | 20 |
| **Distributed Training** | |
| Accelerator | GPU |
| Number of devices | 1 |
| Number of nodes | 1 |
| Number of workers | 24 |
| Precision | 16-bit mixed |
| Strategy | DDP |
